# IDH-mutant gliomas arise from glial progenitor cells harboring the initial driver mutation

**DOI:** 10.1101/2024.10.17.618976

**Authors:** Jung Won Park, Jiehoon Kwak, Keon-Woo Kim, Saehoon Jung, Chang Hyun Nam, Hyun Jung Kim, Sang Mee Lee, Ji-Hyung Park, Jihwan Yoo, Jin-Kyoung Shim, Chungyeul Kim, Sangjeong Ahn, Stefan Pusch, Andreas von Deimling, Jong Hee Chang, Se Hoon Kim, Young Seok Ju, Seok-Gu Kang, Jeong Ho Lee

## Abstract

Discovering the cell-of-origin harboring the initial driver mutation provides a fundamental basis for understanding tumor evolution and development of new treatments. For isocitrate dehydrogenase (IDH)-mutant gliomas – the most common malignant primary brain tumors in adults under 50 – the cell-of-origin remains poorly understood. Here, using patient brain tissues and genome-edited mice, we identified glial progenitor cells (GPCs), including oligodendrocyte progenitor cells (OPCs), as the glioma-originating cell type harboring the IDH mutation as the initial driver mutation. We conducted comprehensive deep sequencing, including droplet digital PCR and deep panel and amplicon sequencing to 128 tissues from 62 patients (29 IDH-mutant gliomas and 33 IDH-negative controls) comprising tumors, normal cortex or normal subventricular zone (SVZ), and blood. Surprisingly, low-level IDH mutation was found in the normal cortex away from the tumor in 38.5% (10 of 26) of IDH-mutant glioma patients, whereas no IDH mutation was detected in the normal SVZ. Furthermore, by analyzing cell-type–specific mutations, the direction of clonal evolution, the single-cell transcriptome from patient brains and novel mouse model of IDH-mutant glioma arising from mutation-carrying OPCs, we determined that GPCs, including OPCs, harboring the initial driver mutation are responsible for the development and evolution of IDH-mutant gliomas. In summary, our results demonstrate that GPCs containing the IDH mutation are the cells-of-origin harboring the initial driver mutation in IDH-mutant gliomas.

## Introduction

IDH-mutant gliomas are the most common brain malignancy in adults under the age of 50, accounting for 12% of all gliomas(*1*). There are two distinct subtypes of IDH-mutant glioma: oligodendroglioma with 1p/19q-codeleted (OD) and astrocytoma with 1p/19q-non-codeleted (AS)(*2, 3*). The majority of *IDH* mutations are heterozygous *R132H* point mutation in *IDH1* (*IDH1^R132H^*), which produces the oncometabolite, D-2-hydroxyglutarate (D-2-HG), implicated in tumorigenesis(*1, 4–6*). Previous single-cell RNA sequencing (scRNA-seq) and bulk RNA sequencing analyses of IDH-mutant gliomas revealed shared glial lineages and developmental hierarchies among AS and OD, suggesting a common progenitor for all IDH-mutant gliomas(*7, 8*). The *IDH* mutation is thought to be the early genetic event in IDH-mutant glioma, a supposition with significant implications for glioma progression(*9–12*). However, whether the *IDH* mutation is indeed the first genetic driver event in IDH-mutant glioma and which cell type in the brain is the common progenitor that acts as the cell-of-origin harboring the *IDH* mutation and drives gliomagenesis remain open questions.

Compared with non-dividing mature cells, stem cells or progenitor cells, characterized by their proliferative capacity, are more likely to acquire somatic mutations owing to intrinsic errors in DNA replication during development and aging(*13, 14*). In the brain, neural stem cells (NSCs), glial progenitor cells (GPCs), including oligodendrocyte progenitor cells (OPC), or microglia that display such proliferative capacity can be a source of the initial somatic mutation that clonally expands and leads to various neurological disorders, such as epilepsy, brain tumors, and neurodegenerative disorders(*15–21*). In line with this, we and others have previously shown using patients’ tissues and mouse models that NSCs located in the subventricular zone (SVZ) are the cells-of-origin harboring the initial driver mutations in IDH-wildtype glioblastoma (GBM) and BRAF-mutant glioma with epilepsy(*22–24*). However, the cell-of-origin harboring the initial driver mutation in IDH-mutant glioma patients has yet to be explored.

To probe the cell-of-origin harboring the initiating *IDH* mutation in glioma patients, we here performed comprehensive deep sequencing procedures, including droplet digital PCR and deep panel and amplicon sequencing, on 128 tissues from 62 individuals – 29 with IDH-mutant gliomas and 33 with IDH-negative controls) – comprising tumors, normal cortex or SVZ tissues, and blood. Interestingly, a low level of *IDH* mutation was observed in normal cortex away from the tumor, but not in normal SVZ tissues (Fig. 1A). We further analyzed cell-type–specific mutations and the direction of clonal evolution in patient brain tissues, discovering that GPCs, including OPCs, in the normal cortex contained the *IDH* mutation as the initial driver mutation. Moreover, by introducing the *IDH* mutation and relevant driver mutations into OPCs, we were able to generate a mouse IDH-mutant glioma model that recapitulated the pathological and molecular features of human IDH-mutant gliomas and their evolution (Fig. 1A). Our study shows that GPCs containing the *IDH* mutation are the cells-of-origin harboring the initial driver mutation in IDH-mutant gliomas.

**Fig. 1.**
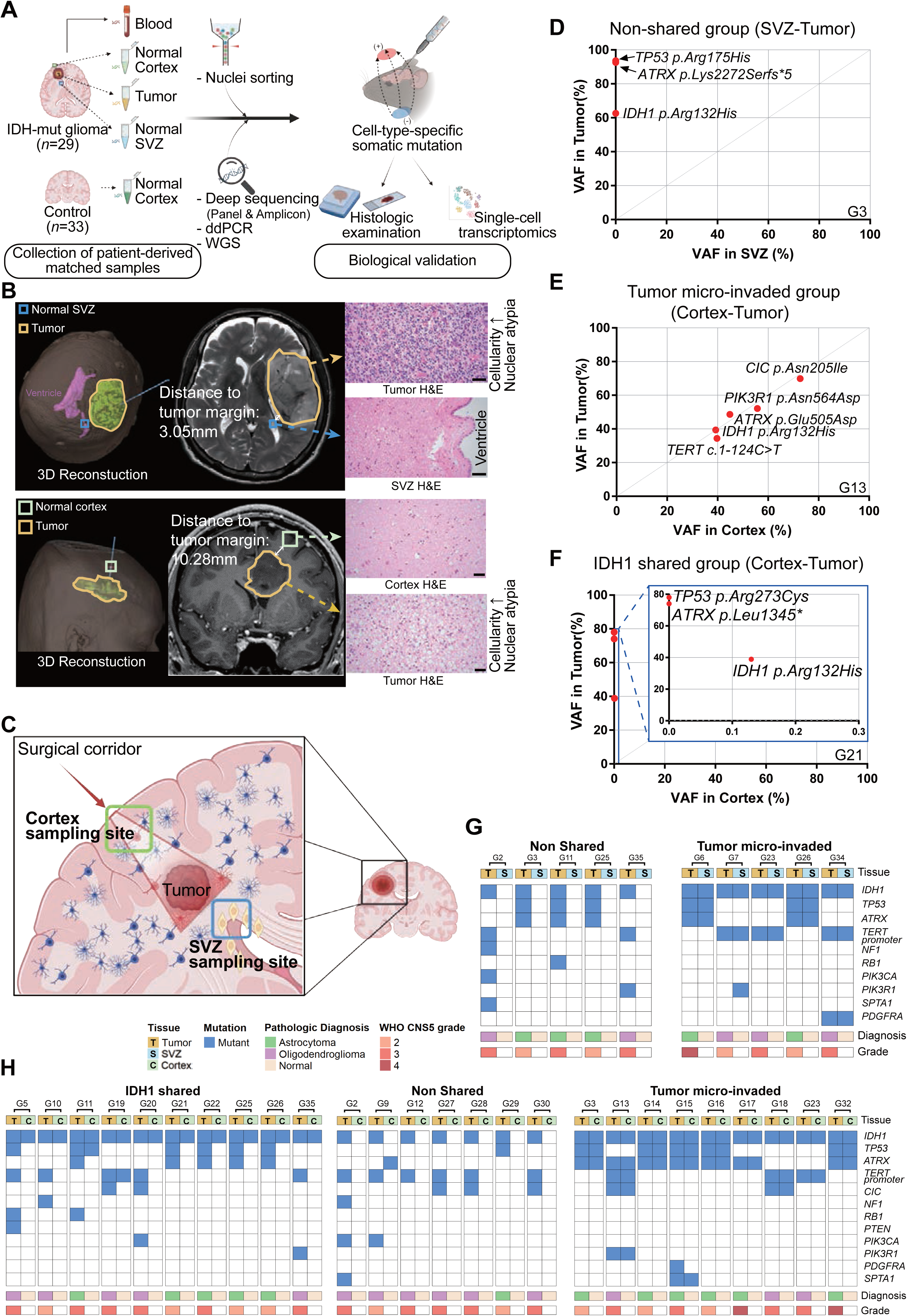
Normal cortex of IDH-mutant glioma patient harbors low-level *IDH* mutation as the initial driver mutation. (**A**) Schematic depiction of the experimental design of the study showing the comprehensive deep sequencing analysis of tumor and matched normal brain tissues from IDH-mutant glioma and control individuals, followed by biological validation in mouse IDH-mutant glioma models. Created with BioRender.com. (**B**) Representative 3-dimensional MR images showing sampling site and distance to tumor margin and an H&E examination of histologic features. Scale bars, 50 μm. Representative case of normal SVZ and tumor (top) and normal cortex and tumor (bottom). (**C**) Illustration of the neurosurgical procedure, with the surgical corridor for accessing the normal cortex, SVZ, and tumor tissues of IDH-mutant glioma individuals. Blue cells, normal glial cells; red cells, mutant glial cells; yellow cells, NSCs. Created with BioRender.com. (**D** to **F**) Representative VAF scatterplots of somatic mutations from matched samples. Red dots indicate driver mutations found in each sample. Mutations restricted to the SVZ (or cortex; *x*-axis), the tumor (*y*-axis), or shared between the two tissues are shown (see Figs. S3 and S4). (D) Representative VAF scatterplot of somatic mutations in the SVZ and tumor of the Non-shared group (G3 individual). (E) Representative VAF scatterplot of somatic mutations in the cortex and tumor of the Tumor micro-invaded group (G13 individual). (F) Representative VAF scatterplot of somatic mutations in the cortex and tumor of the IDH1-shared group (G21 individual). fs, frameshift. *, termination. (**G** and **H**) Summary of SVZ–tumor (G) and cortex–tumor (H) relationship. Matched specimens from the same individuals are marked by square brackets (‘┌┐’) at the tops of columns.

## Results

### Low-level IDH1^R132H^ presents as the initial driver mutation in the normal cortex of IDH-mutant glioma tissues

To explore the initial driver mutation in IDH-mutant gliomas, we inferred the relative timing of somatic driver events by reconstructing them using whole-genome sequencing (WGS) data from 38 IDH-mutant glioma patients (20 from the PCWAG database, 12 from TCGA, 6 from our cohort)(*3, 25, 26*). Aggregating the ordering information about the timing of each driver point mutation, as well as the timing status of clonal and subclonal copy number segments across samples, we defined a probabilistic ranking of driver events that represents the relative timing of driver events over tumor evolution(*26*). This analysis revealed that the *IDH1* mutation is the earliest event, followed by other driver mutations, such as loss of 1p/19q, *TERT* promoter mutations, *TP53*, and *ATRX* (fig. S1A). This finding aligns with previous studies showing that *IDH* mutations were the only mutation in common among all recurrent tumors from IDH-mutant glioma patients, and that there were no cases where an *IDH1* mutation occurred after *TP53* mutation or 1p/19q loss(*10–12*). Our results suggest that the *IDH1* mutation is the initial driver in IDH-mutant glioma.

To probe the origin of the *IDH* mutation as the initial driver mutation, we examined whether *IDH* mutations or other driver mutations were present in the normal SVZ or cortex containing NSCs or GPCs (including OPCs) in IDH-mutant glioma patients. We obtained 95 triple- or quadruple-matched samples that included normal SVZ or cortex (confirmed pathologically and radiologically), tumor tissue, and blood from 29 individuals with AS or OD who underwent supra-total resection surgery (Fig. 1A to C, and table S1). For controls, we collected 33 pathologically normal cortex samples from IDH-negative individuals diagnosed with epileptic diseases, Alzheimer’s disease, autism spectrum disorder, metastatic brain tumors, IDH-wildtype glioma, or other non-GBM conditions (table S2). Previous studies reported low-level mosaicism of the *TERT* promoter in the normal SVZ of IDH-wildtype GBM or *IDH* mutation in the cortex of non-neoplastic individual(*22, 27*) To accurately detect low-level somatic driver mutations in the normal SVZ or cortex of IDH-mutant glioma patients, we employed droplet digital PCR (ddPCR) of the *IDH1^R132H^*mutation, deep sequencing of glioma-related genes (average read depth, 757.5×), and ultra-deep targeted amplicon sequencing (average read depth, 391,958×). Using ddPCR of *IDH1^R132H^* in 33 normal cortex samples from *IDH* mutation-negative individuals, we determined a variant allele frequency (VAF) cut-off for true detection of an *IDH1* mutation of 0.115%, a value three standard deviations from the mean of noise or artifactual VAF (*P* < 0.00135), and applied it to all samples (See Materials and Methods and fig. S1B). For deep-panel sequencing of 15 glioma-related genes in all samples (table 3), we considered overlapping somatic mutation calls in Mutect2 and Strelka2 as true somatic mutations (see Materials and Methods). These were validated using ultra-deep targeted amplicon sequencing, taking into account somatic mutations with statistically significant VAFs (*P* < 0.05, compared with background errors) as true somatic mutations(*28*) (see Materials and Methods).

On the basis of this analysis, we were able to classify IDH-mutant glioma patients into three groups based on mutations observed in the SVZ and cortex: (i) a Non-shared group, with no somatic mutations shared with tumors; (ii) a Tumor micro-invaded group, where all somatic mutations, including driver mutations, are shared with tumors; and (iii) an IDH1-shared group, in which the normal SVZ or cortex shares only the *IDH1^R132H^*or *IDH1^R132H^* plus one additional early driver event with tumors (Fig. 1D to F, fig. S3 to S5, and table S4 and S5). There were no significant differences among these three groups with respect to the distance to tumor margin or tumor grade (fig. S1C and D). An analysis of the normal SVZ classified IDH-mutant glioma patients into two groups – Non-shared and Tumor micro-invaded – each of which accounted for 50% (5 of 10) of the total (Fig. 1G). Notably, there were no patients in the IDH1-shared group. In contrast, in the normal cortex, 38.5% (10 of 26) of IDH-mutant glioma patients were classified into the IDH1-shared group (Fig. 1H), primarily carrying only the *IDH1* mutation, with two cases also having an early driver event, exemplified by a *TP53* mutation in G11 and a *TERT* promoter mutation in G19. Moreover, both AS and OD were similarly observed in the IDH1-shared group of cortices (Fig. 1H). Since OD is defined by an *IDH* mutation and 1p/19q-codeletion, we further investigated for the presence of 1p/19q-codeletion in the normal cortex of the IDH1-shared group using a fluorescence *in situ* hybridization (FISH)-based assessment. We found no chromosomal defects with an obviously high ratio (Test probe/Control probe > 1) in the normal cortex from OD patients in the IDH1-shared group (fig. S2A) – similar to the case for normal cortex from AS without 1p/19q-codeletion (fig. S2B) – whereas loss of chromosomes with a low ratio (Test probe/Control probe < 0.8) was clearly observed in the OD tumor (fig. S2C). Consistent with a previous study of multiple patients’ tissues showing no cases in which an *IDH1* mutation occurred after the loss of 1p/19q(*11*), this result suggests that the *IDH* mutation is the initial driver event in OD patients, occurring before the loss of 1p/19q. In the normal cortex, Non-shared and Tumor micro-invaded categories accounted for 26.9% (7 of 26) and 34.6% (9 of 26), respectively, of the total (Fig. 1H). These findings, which contrast with a previous report of early driver events, including *TERT* promoter and *EGFR* mutations, in the normal SVZ of IDH-wildtype GBM(*22*), suggest that the *IDH1^R132H^* mutation in IDH-mutant glioma likely arises from the normal cortex: IDH-mutant gliomas showed a weak association with NSCs (0%) relative to IDH-wildtype GBM accompanied by *TERT* promoter alteration (56.25%). (fig. S3C). Taken together, our results suggest that the initial driver *IDH* mutation event in IDH-mutant glioma originates from the normal cortex, where clones of the cell-of-origin harboring the *IDH* mutation are located.

### GPCs in the cortex are the cells-of-origin harboring the IDH1^R132H^ mutation

To further verify the direction of clonal evolution between the cortex harboring the *IDH1* mutation and the tumor, we examined the normal cortex for the absence of most tumor-restricted mutations, including driver mutations (other than the *IDH* mutation). If these mutations are absent, it would suggest that the initial clone with the *IDH* mutation in the cortex acquires tumor-restricted mutations during clonal evolution into a tumor. To test this prediction, we performed WGS on matched tumor and normal tissues from four IDH-mutant glioma patients of the IDH1-shared group. These patients had either only the *IDH1^R132H^*mutation (G20, G21, G35) or *IDH1^R132H^* plus a *TP53* mutation (an early driver event) (G11) in their normal cortex. We then randomly selected 58 tumor-restricted variants (9-18 variants per individual), including all driver mutations, and examined the normal cortex for their presence using ultra-deep targeted amplicon sequencing (average read depth, 160,395×). As expected, tumor-restricted mutations were rarely observed in the normal cortex across all four cases (Fig. 2A to D). For individual G35, we also performed WGS on both initial and recurrent tumors, cataloging the initial tumor- or recurrent tumor-restricted mutations (Fig. 2B to D). We then conducted ultra-deep targeted amplicon sequencing of these mutations in the cortex, comparing their presence between the cortex, initial tumor, and recurrent tumor. Interestingly, the normal cortex shared no initial or recurrent tumor-restricted mutations other than *IDH1^R132H^* (Fig. 2D). Additionally, most initial tumor-restricted mutations, including *NRAS, PIK3R1*, and *NOTCH1* driver mutations and *CDKN2A/B* homozygous deletion, were not shared with the recurrent tumor. Only three driver mutations – *IDH* and *TERT* promoter mutations and 1p/19q codeletion – plus one tumor-restricted mutation (*HRNR*) were shared between initial and recurrent tumors (Fig. 2C and D). This suggests that both the initial and recurrent tumors were derived from a common ancestor clone harboring *IDH* and *TERT* promoter mutations with 1p/19q loss, originating from an initial clone in the cortex with only the *IDH* mutation (Fig. 2C). Consistent with our findings, previous studies reported that a common ancestor clone harboring the *IDH* mutation contributes to both initial and recurrent IDH-mutant gliomas(*10, 12, 29*). Collectively, our results suggest that cells in the normal cortex harboring the *IDH* mutation clonally evolve into tumor cells by acquiring additional driver mutations.

**Fig. 2.**
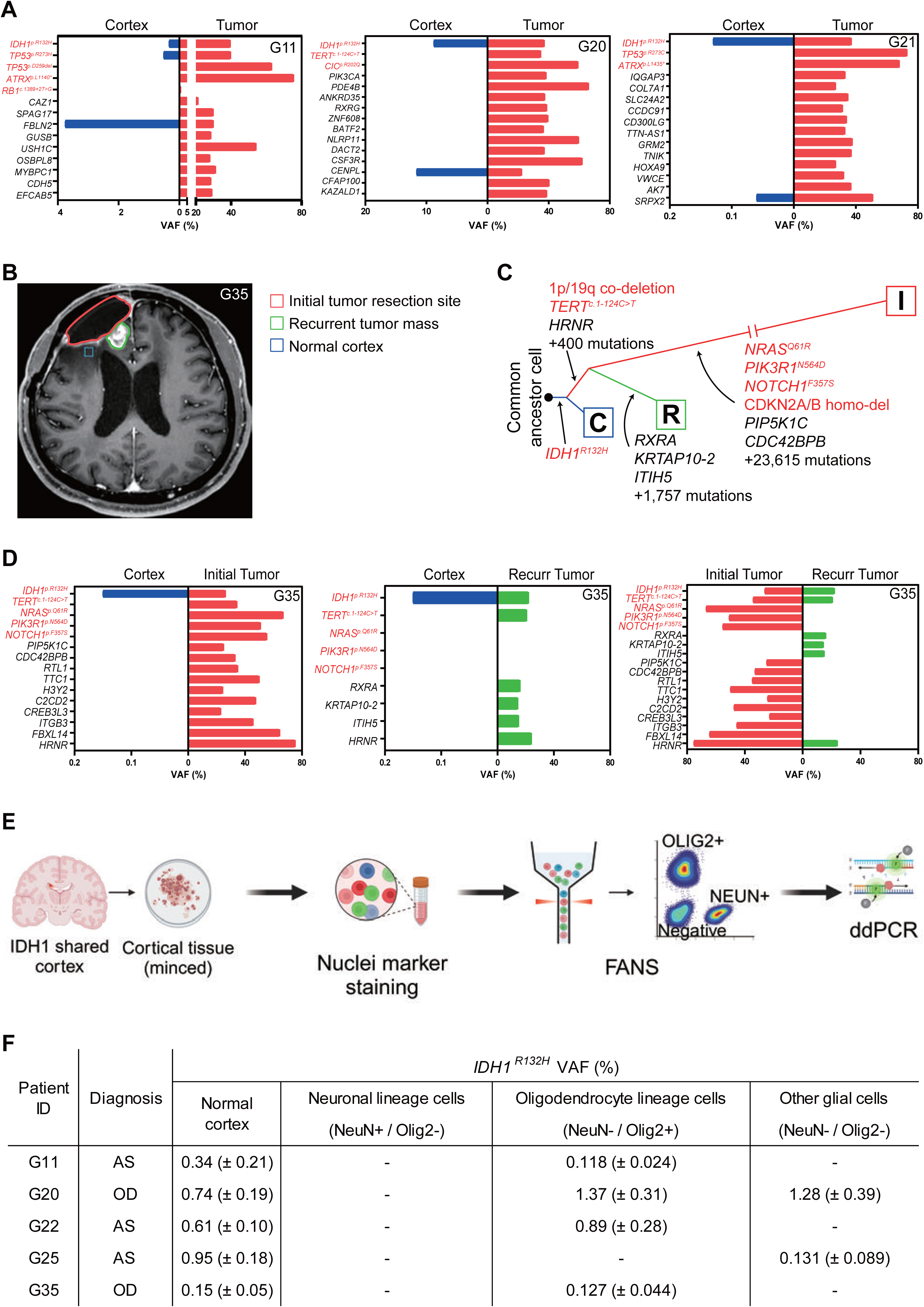
GPCs, including OPCs, are the cells-of-origin harboring the *IDH* mutation. (**A**) Bar graphs showing VAF of tumor-restricted mutations in normal cortex and tumor tissue from three individuals (G11, G20, G21). Red-labeled genes indicate driver mutations. (**B** to **D**) Illustrative longitudinal case (G35 patient) showing the evolutionary relationships within the normal cortex, initial tumor, and recurrent tumor. (B) Representative MR image. Image was acquired at diagnosis of recurrent tumor mass. (C) A phylogenetic tree depicting the patterns of clonal evolution within three tissues adopted by tumor-restricted mutations. C, Normal cortex. I, Initial tumor. R, Recurrent tumor. Red genes, driver mutations. (D) Bar graphs showing VAF of tumor-restricted mutations in normal cortex, initial tumor, and recurrent tumor tissue. Red-labeled genes indicate driver mutations. (**E**) Schematic figure showing deep sequencing analysis of cell-type–specific *IDH* mutation. Nuclei sorting according to cell type was performed, followed by ddPCR analysis for *IDH1^R132H^*. OLIG2, marker for oligodendrocyte-lineage cells; NEUN, marker for neuronal-lineage cells; double-negative populations, other glial cells, including astrocytes. Created with BioRender.com. (**F**) Cell-type–specific distribution of *IDH1^R132H^* (identified in bulk normal cortex) in the normal cortex from five IDH-mutant glioma individuals. ‘-‘ denotes mutation negative. OD: oligodendroglioma, IDH-mutant, 1p/19q-codeleted; AS: astrocytoma, IDH-mutant, 1p/19q-non-codeleted.

To identify the specific brain cell-type in the cortex that carries the *IDH* mutation and serves as the cell-of-origin for IDH-mutant gliomas, we sorted cortical nuclei labeled with cell-type– specific markers such as, NEUN and OLIG2, from five IDH-mutant glioma individuals in the IDH1-shared group (Fig. 2E and fig. S6). We isolated three types of nuclei: (i) NEUN-positive and OLIG2-negative, corresponding to neurons; (ii) NEUN-negative and OLIG2-positive, corresponding to oligodendrocyte-lineage cells, including OPCs and mature oligodendrocytes; and (iii) NEUN-negative and OLIG2-negative cells, corresponding to “other” glial-lineage cells, including astrocytes. On average, we obtained 77,041 neuronal nuclei, 257,960 oligodendrocyte-lineage nuclei, and 42,723 other glial-lineage nuclei per cortex. Isolated nuclei, as well as their corresponding bulk normal cortex, were simultaneously subjected to ddPCR analysis for *IDH1^R132H^* (Fig. 2F). Again, the bulk of the normal cortex was positive for *IDH1^R132H^*. Surprisingly, no neurons from any individual were positive for the *IDH* mutation (Fig. 2F). However, oligodendrocyte-lineage cells were *IDH* mutation-positive positive in 4 of 5 individuals, whereas “other” glial-lineage cells, which includes astrocytes, were positive in 2 of 5 individuals (Fig. 2F). Notably, both oligodendrocytes-and other glial-lineage cells were positive for the *IDH* mutation in individual G20 (Fig. 2F). Given that GPCs can give rise to OPCs, oligodendrocytes and astrocytes in human brain development(*30*), our results suggest that GPCs, including OPCs, in the cortex are the cells-of-origin harboring the *IDH* mutation. Their progeny cells likely acquire additional driver mutations over time, ultimately evolving into IDH-mutant gliomas.

### Development of a mouse model of OPC-originating IDH-mutant glioma

We next sought to verify that GPCs or OPCs harboring an *IDH* mutation proliferate and ultimately evolve into IDH-mutant gliomas *in vivo*. Given that GPC-specific markers are less completely characterized, we utilized the well-defined OPC-specific marker, Ng2, to develop a PiggyBac transposon system incorporating *Ng2* enhancer-driven Cre (Ng2-Cre)(*31*) (fig. S7A) and single-guide RNAs (sgRNA) targeting glioma driver genes or LacZ (control) into the genome of an *Idh1^LoxP(R132H)/+^;*loxP-stop-loxP (LSL)-Cas9-EGFP^fl/+^ mouse. Postnatal electroporation of these PiggyBac plasmids into the right lateral ventricle induced *Idh1^R132H^*or glioma driver mutations in a small fraction of OPCs, which are derived from neural stem cells carrying PiggyBac plasmids and are tracked by a co-expressed EGFP reporter (Fig. 3A). In this system, we validated that GFP-positivity co-localized with OPC and oligodendrocyte-lineage markers (Ng2 and Olig2), but not astrocyte (Gfap) or neuronal (NeuN) markers (fig. S7B to E). Sanger or amplicon sequencing showed that GFP-positive cells exhibited heterozygous *Idh1^R132H^* and CRISPR/Cas9-targeted driver mutations (fig. S8A to C)(*32, 33*). Introducing an *Idh1^R132H^*mutation alone into OPCs increased the number of GFP-positive cells, suggesting enhanced proliferation of *Idh1*-mutation–expressing OPCs in the OPC-I (*Idh1^R132H^*) mouse (Fig. 3B and fig. S9A). Interestingly, and in line with a previous report identifying a focal cluster of *IDH*-mutant clones in postmortem normal white matter(*27*), we similarly found focal clusters of GFP-expressing *Idh1*-mutant clones in mouse normal white matter (fig. S9B). Previous studies showed that a single *Idh* mutation in neural stem cells results in increased proliferation but is insufficient to induce tumor formation(*34, 35*). Consistent with these previous studies, an *Idh* mutation alone did not induce tumor formation in OPC-I mice (fig. S9C).

**Fig. 3.**
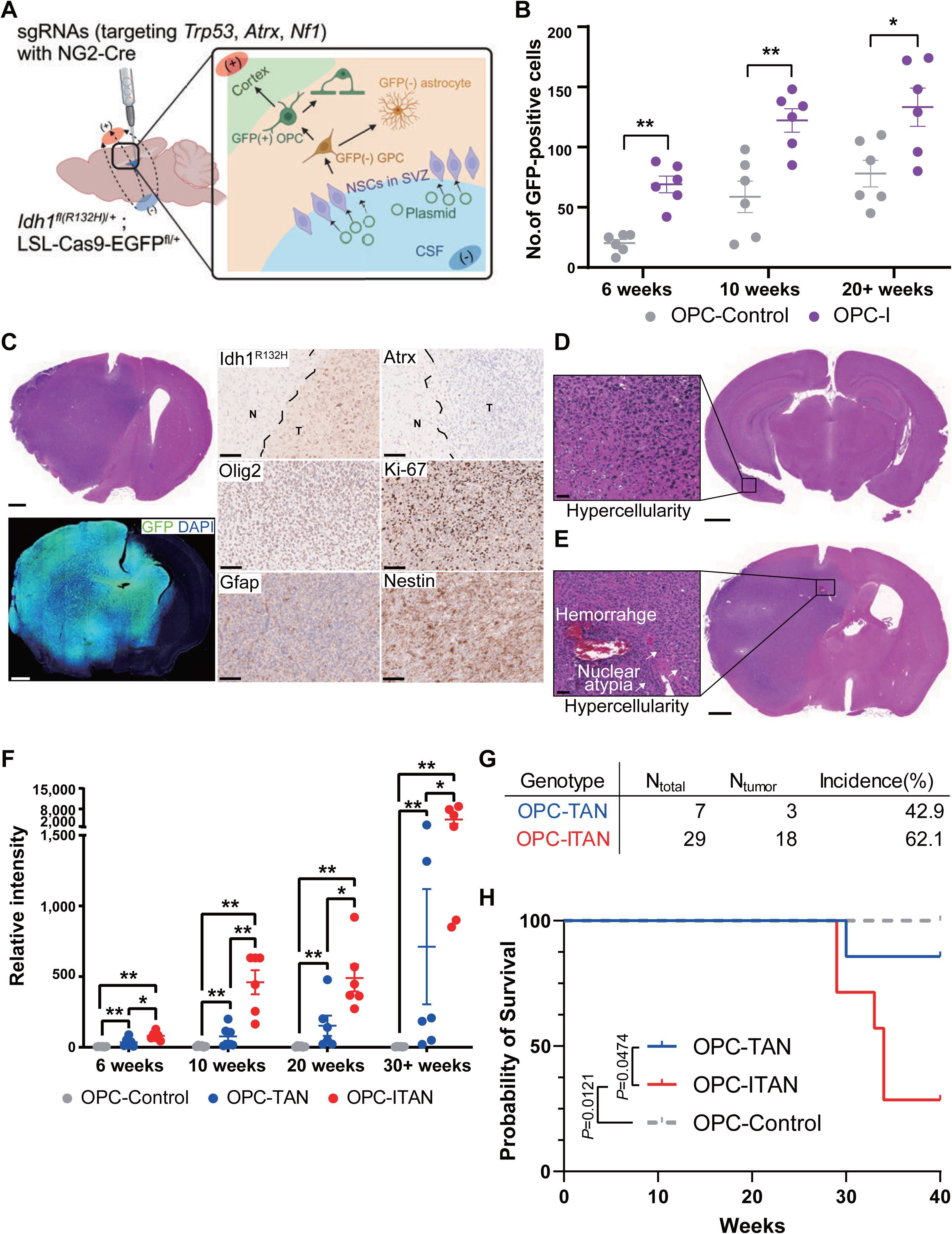
Development of a mouse model of IDH-mutant glioma arising from OPCs. (**A**) Experimental scheme showing *in vivo* electroporation of PiggyBac vectors containing sgRNAs against *Trp53, Atrx, Nf1, Cdkn2a,* and *LacZ* (control) under the control of Ng2-Cre, which induces the OPC-specific expression of Cre recombinase and knockout of target genes in *Idh1^fl(R132H)/+^*;LSL-Cas9-EGFP^fl/+^ mice at postnatal day 0-1 (P0-1). Oligodendrocyte-lineage cells (including OPCs) are GFP-positive, whereas neurons and astrocytes were GFP-negative (see fig. S7). CSF, cerebrospinal fluid in the lateral ventricle. Created with BioRender.com. (**B**) Scatter dot-plot showing the number of GFP-positive cells in the OPC-control (*Idh1^wildtype^*) and OPC-I (*Idh1^R132H^*) mouse. The sections selected from each mouse brain were those that included the highest number of GFP-positive cells. Data are presented as means ± s.e.m. (error bars); *n* = 6 per time point. \**P* < 0.05, \*\**P* < 0.01 (Student’s two-tailed *t*-test). (**C**) Representative images of the OPC-ITAN (*Idh1^R132H^*;*Trp53*;*Atrx*;*Nf1*) model mouse. Top left: H&E-stained image of the whole brain; bottom left; immunofluorescence image of the whole brain; center and right: immunohistochemical staining for the IDH-mutant glioma markers, Idh1^R132H^, Atrx, Olig2, Ki-67, Gfap, and Nestin. T, tumor area; N, normal area. The sections selected from each mouse brain were those that showed the most malignant lesions. Scale bars, 1,000 μm (left) and 100 μm (center and right). (**D** and **E**) Representative histology slides of IDH-mutant glioma model mouse showing different tumor grades. Insets indicate the malignant lesion in each mouse brain with corresponding histopathological findings. Scale bars, 50 μm (left) and 1,000 μm (right). (D) H&E images from an OPC-ITAN model mouse (34 weeks old) with low-grade glioma. Left: Magnified image of hypercellularity area; right: whole brain image. (E) H&E images of an OPC-ITAN model mouse (29 weeks old) with high-grade glioma. Left: Magnified image of hypercellularity, nuclear atypia and intra-tumoral hemorrhage area; right: whole brain image. Arrows, nuclear atypia. (**F**) Scatter dot-plot showing quantification of the relative intensity of GFP-positive signals in OPC-TAN (*Idh1^wildtype^*;*Trp53*;*Atrx*;*Nf1*) and OPC-ITAN (*Idh1^R132H^*;*Trp53*;*Atrx*;*Nf1*) mouse models, compared with the intensity of GFP-positive signals in OPC-control (*Idh1^wildtype^*;LSL-Cas9-EGFP^fl/+^) mice at post-electroporation day 7. The sections selected from each mouse brain were those that included the highest GFP intensity sites. Data are presented as means ± s.e.m. (error bars); *n* = 6 per time point. \**P* < 0.05, \*\**P* < 0.01 (Student’s two-tailed *t*-test). (**G**) Summary data showing tumor incidence up to 40 weeks of age in OPC-TAN and OPC-ITAN mouse models. Low-grade gliomas without any obvious external symptoms were diagnosed after harvesting the brain at 40 weeks. N, number of mice in each genotype. (**H**) Kaplan-Meier survival graph for OPC-control (*n* = 6), OPC-TAN (*n* = 7), and OPC-ITAN (*n* = 7) model mice (log-rank test).

We then examined the role of additional glioma driver mutations in tumor formation by combining the *Idh* mutation with deletions of the transformation-related protein 53 gene (*Trp53*; also known as tumor suppressor protein 53 [p53]), alpha-thalassemia/mental retardation syndrome X-linked gene (*Atrx)*, neurofibromin 1 gene (*Nf1*) and cyclin-dependent kinase inhibitor 2a gene (*Cdkn2a)*, which were recurrently identified mutations in human IDH-mutant gliomas as well as our patient cohort. It was previously reported that concurrent mutations in *Idh1, Trp53*, and *Atrx* genes in brain cells of the mouse were insufficient to induce tumor formation(*36*). In accord with this, OPC-ITAC mice, with concurrent mutations in *Idh1, <u>T</u>rp53, <u>A</u>trx*, and *<u>C</u>dkn2a*, showed no evidence of tumor formation (fig. S10A to C). However, concurrent *Idh1, <u>T</u>rp53, <u>A</u>trx*, and *<u>N</u>f1* mutations (OPC-ITAN mice) led to formation of IDH-mutant astrocytomas, as confirmed by positive immunostaining for Olig2, Ki67, Nestin and GFAP, and negative staining for Atrx (Fig. 3C). Our OPC-ITAN model mice exhibited histological features ranging from low-grade gliomas with hypercellularity to high-grade gliomas with nuclear atypia and intratumoral hemorrhage (Fig. 3D and E). These phenotypes align well with human data showing varying tumor grades associated with *IDH* mutations(*37*). We then examined the effect of *Idh* mutation on tumor formation in combination with *<u>T</u>rp53, <u>A</u>trx*, and *<u>N</u>f1* mutations by generating an OPC-TAN (*Idh1^wildtype^*;*Trp53*;*Atrx*;*Nf1)* mouse. Tumor cell proliferation and tumor formation were decreased in the OPC-TAN mouse compared with the OPC-ITAN mouse, but the former showed better survival (Fig. 3F to H, and fig. S11), suggesting that the *IDH* mutation combined with other glioma driver mutations results in formation of more aggressive tumors. To exclude the possibility of tumor formation from mutation-acquiring mature cortical cells, such as neurons, astrocytes and oligodendrocytes, we introduced *Idh1, Trp53, Atrx*, and *Nf1* mutations into differentiated cortical cells of *Idh1^fl(R132H)/+^*;LSL-Cas9-EGFP^fl/+^ mice using AAV5-Cre (fig. S12A to E). These mice showed no significant cell proliferation or tumor formation (fig. S12F and G). Taken together, data from our mouse IDH-mutant glioma models demonstrate that OPCs harboring the *IDH* mutation proliferate, acquire additional glioma driver mutations, and ultimately evolve into IDH-mutant gliomas.

### A mouse model of IDH-mutant glioma effectively recapitulates human IDH-mutant glioma at the single-cell transcriptome level

To further determine whether our mouse IDH-mutant glioma model accurately represents human IDH-mutant glioma at the single-cell transcriptome level, we performed scRNA-seq on four OPC-ITAN mice (26-38 weeks old) and one control mouse (20 weeks old) (fig. S13A). A total of 21,828 of GFP-positive and -negative cells were analyzed from these five mice brains and cell-type annotations were assigned to each cluster based on established methods(*7, 8, 38*). This marker gene-based classification identified major normal cell types, including oligodendrocytes (OL), committed oligodendrocyte precursor cells (COP), astrocytes (AC), neural stem cells (NSC), neurons, microglia (MG), ependymal cells (Epen), endothelial cells (Endo), pericytes (PC), and T-cells (Fig. 4A, fig. S13B). After labeling normal cells, we distinguished tumor cells from OPCs using the tumor marker genes, *Cas9-EGFP, Mki67*, and *Nes*(*39, 40*) (Fig. 4A, fig. S13B). Because we introduced driver mutations into OPCs with co-expression of the GFP reporter, we defined cell types expressing GFP – which constituted >30% of the population – as ‘tumor-lineage cells’, which included OPCs, COPs, and tumor cells (fig. S14A to C).

**Fig. 4.**
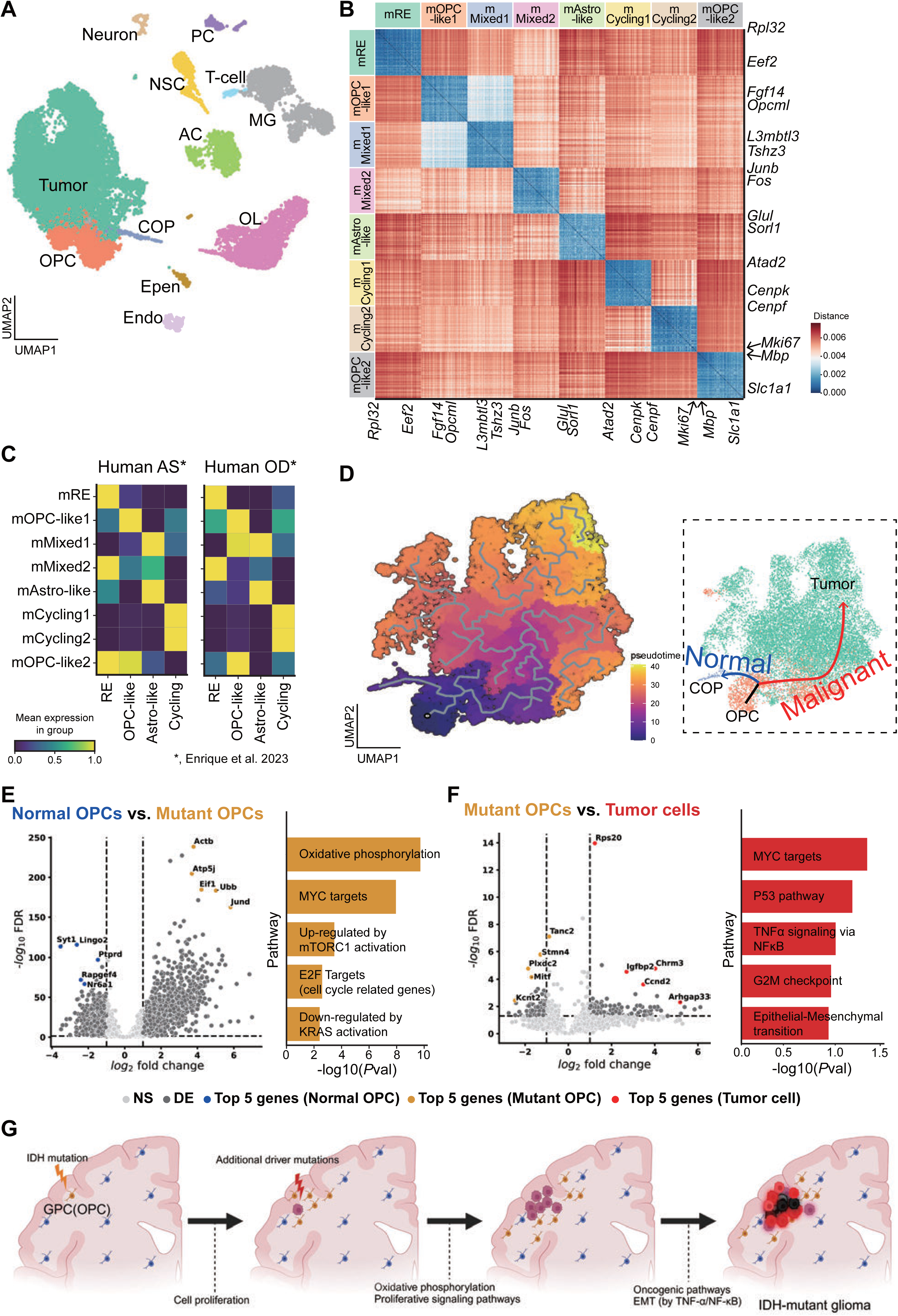
Transcriptomic landscape of mouse IDH-mutant glioma model recapitulating human IDH-mutant gliomas at single-cell resolution. (**A**) UMAP representation of major cell-type clusters within brains from four IDH-mutant glioma model (OPC-ITAN) mice and a control (OPC-control) mouse. Colors indicate cell-type assignments. (**B**) Correlation plot showing the clustering of mouse IDH-mutant glioma-associated transcriptomic modules. Modules were identified by NMF-based subclustering analysis of tumor-lineage cells. Each module was termed according to known human IDH-mutant glioma subpopulation markers(*7, 8, 38*). Two representative genes in each module were annotated. (**C**) Matrix plots indicating the expression level of mouse IDH-mutant glioma-associated transcriptomic modules within subpopulations of human IDH-mutant gliomas(*38*). AS, astrocytoma (IDH-mutant, 1p/19q–non-codeleted); OD, oligodendroglioma (IDH-mutant, 1p/19q-codeleted). (**D**) Left: Developmental trajectory of tumor cells based on a pseudotime analysis, illustrated as a UMAP representation of tumor-lineage cells. OPC – the cell-of-origin in our mouse IDH-mutant glioma model – is the root cell (marked with a black dot). Right: Two distinct developmental paths, depicted as a UMAP representation of tumor-lineage cells with labels. (**E** and **F**) Volcano plots of differentially expressed gene (DEG) analysis with GSEA using the Hallmark gene set(*70*). The top five upregulated genes are labeled blue in normal OPCs, orange in mutant OPCs, and red in tumor cells OPCs. *P*-value threshold = 0.05; log2 fold change threshold = 1. NS, not significant. DE, differently expressed gene. (E) Comparison of mutant OPCs with normal OPCs(*44*). (F) Comparison of tumor cells with mutant OPCs. (**G**) Illustration showing the developmental process in IDH-mutant gliomas. EMT, epithelial-mesenchymal transition. Blue cells, normal GPCs (OPCs). Created with BioRender.com.

We then identified distinguishable transcriptomic features of our mouse model using unbiased, non-negative matrix factorization (NMF) of tumor-lineage cells. NMF is a widely used analytical tool for revealing the features – even sparse properties – of high-dimensional data in a given dataset(*38, 41, 42*). Our NMF analysis sorted mouse IDH-mutant glioma-associated transcriptomics into eight distinct modules (fig. S15A and B). Based on known human IDH-mutant glioma subpopulation gene sets(*7, 8, 38*), we labeled these modules as follows: 1) mouse ribosome-enriched (mRE) (*Rpl32, Eef2*); 2) mouse OPC (mOPC)-like 1 (*Fgf14, Opcml*); 3) mOPC-like 2 (*Mbp, Slc1a1*); 4) mouse astrocyte (mAstro)-like (*Glul, Sorl1*); 5) mouse cycling cell 1 (mCycling1) (*Cenpk, Atad2*); 6) mCycling2 (*Cenpf, Mki67*); 7) mouse mixed 1 (mMixed1) (*L3mbtl3, Tshz3*); and 8) mMixed2 (*Junb, Fos*) (Fig. 4B, fig. S15A to C, and table S9). A comparison of these mouse glioma-associated modules with published single-nucleus RNA-seq data for human IDH-mutant glioma tumor cells(*38*) revealed a remarkable match in all modules, except for the two mMixed programs, which showed mixed expression patterns within AS and OD (Fig. 4C). Hierarchical clustering of pseudobulk tumor cell transcriptomic data from our mouse model, human IDH-mutant gliomas(*38*), and human IDH-wildtype GBM(*43*) demonstrated that the mRNA profile of our mouse model was more similar to that of human IDH-mutant gliomas than human IDH-wildtype GBM (fig. S16). These findings suggest that our mouse IDH-mutant glioma model replicates the transcriptomic features of human IDH-mutant gliomas at single-cell resolution.

We further reconstructed the developmental pathway of tumor-lineage cells from OPC to tumor cell using the scRNA-seq dataset (Fig. 4D). This trajectory analysis revealed two separate developmental paths: a normal trajectory from OPC to COP, and a malignant trajectory from OPC to tumor cell (Fig. 4D). A pseudotime analysis showed that different developmental stages employed distinct glioma-associated modules (fig. S17). For instance, the mMixed2 and mRE modules were active during the transition from OPC to tumor cells and early malignant stages, whereas the mCycling1 and mCycling2 modules were involved in late malignant stages. In contrast, the mOPC-like2 module predominantly operated in the normal developmental trajectory. To understand how OPCs evolve into tumor cells after acquiring driver mutations, we performed a differential gene expression analysis followed by gene set enrichment analysis (GSEA) of normal OPCs, mutant OPCs, and tumor cells. Because of the limited number of GFP-negative normal OPCs in our dataset, we compared scRNA-seq data for GFP-positive OPCs (mutant OPCs) from our mouse models with published data for adult mouse OPCs(*44*). Our analysis showed that oxidative phosphorylation and proliferative signaling pathways (e.g., Myc, mTOR, and Kras pathways) were enriched in mutant OPCs compared with normal OPCs (Fig. 4E). The top upregulated genes in mutant OPCs included the β-actin gene, *Actb* (generally deregulated in the majority of tumor cells(*45, 46*)); the ATP synthesis-related gene, *Atp5j* (increased expression in breast cancer(*47*)); the translation-regulatory factor gene, *Eif1* (upstream regulator of the translation initiation step of protein synthesis employing the PI3K/AKT/mTOR pathway(*48*)); one of the *Jun* transcription factor components, *Jund* (a crucial substrate of JNK signaling and RAS-driven tumorigenesis in many tissues(*49*)); and the protein turnover-related gene, *Ubb* (independent prognostic, predictive marker for glioma(*50*)). In tumor cells, we observed upregulation of the oncogenic signaling pathways Myc and P53, and the TNF-α/NF-κB–accelerated epithelial-mesenchymal transition program compared with mutant OPCs (Fig. 4F). The top upregulated genes in tumor cells included *Rps20* (ribosomal protein), a marker of poor prognosis in IDH-wildtype GBM(*51*); *Igfbp2*, a cell proliferation modulator in IDH-wildtype GBM(*52, 53*); and *Ccnd2*, a regulator of cell cycle proteins(*54*). Overall, our mouse IDH-mutant glioma model effectively replicates human IDH-mutant gliomas, illustrating the molecular evolution from OPCs (or GPCs), through acquisition of *IDH* mutations and subsequent driver mutations, to tumor cells of IDH-mutant gliomas (Fig. 4G).

## Discussion

Here, we analyzed 128 tissues from 62 individuals comprising tumors (including one recurred tumor), normal cortex or SVZ, and blood, in conjunction with a novel mouse model of IDH-mutant glioma to identify GPCs, including OPCs, as the cells-of-origin harboring the initial *IDH* driver mutation. A comprehensive deep sequencing analysis of low-level *IDH* mutations, aided by analyses of cell-type–specific mutations and the direction of clonal evolution in patients’ tissues, revealed that GPCs, including OPCs, in normal cortex harbored an *IDH1^R132H^* mutation as the initial driver mutation and genetically evolved into IDH-mutant glioma in 38.5% (10 of 26) of IDH-mutant glioma patients. We further developed an OPC-ITAN mouse model of IDH-mutant glioma arising from OPCs that recapitulated the histopathologic and transcriptomic features as well as molecular evolution of human IDH-mutant glioma. Thus, our study provides direct evidence that IDH-mutant glioma arises from GPCs (including OPCs) harboring an *IDH* mutation as the initial driver mutation.

Recent scRNA-seq studies of IDH-mutant glioma patients suggest that two distinct subtypes of IDH-mutant glioma – AS and OD – originate from the same glial lineage and share developmental hierarchies from a common cell-of-origin(*7, 8*). A study that included serial biopsies from 321 gliomas found that *IDH1* mutations never occurred after *TP53* mutations or 1p/19q loss, indicating that *IDH1* mutations are the earliest genetic event(*11*). IDH-mutant GBM (now classified as IDH-mutant grade 4 astrocytoma) and IDH-wildtype GBM show a similar histopathology, but appear to represent distinct disease entities that originate from a separate cell-of-origin(*55*). However, which brain cells are the cell-of-origin harboring the *IDH* mutation has remained unclear. Our study identified GPCs (including OPCs) in the normal cortex as the common cells-of-origin for IDH-mutant gliomas. These cells acquire *IDH* mutations as an initial driver event and can further evolve into AS or OD through additional mutations, such as mutation of *TP53* and *ATRX* and loss of 1p/19q and. Interestingly, IDH-mutant gliomas predominantly occur in the frontal lobe, in contrast to the more varied locations of IDH-wildtype GBMs(*1, 55, 56*). This frontal lobe predominance may be attributable to the abundance of OPCs in this brain region and their lifelong dividing capacity, which allows them to potentially accumulate mutations(*30, 57–60*).

Recent studies have reported that clones with cancer driver mutations expand in normal tissues as somatic mosaicism(*61, 62*). This phenomenon has been observed in various tissues, including the liver, esophagus, blood, and brain(*27, 63–65*). The association of positively selected clones with common driver mutations suggests their relation to the cells-of-origin of tumors and provides insight into early genetic events in tumorigenesis(*66*). In addition, such clones carrying driver mutations in normal tissues are often predisposed to tumor evolution(*64*). In the brain, clonal oncogenic mutations like *IDH1^R132H^* have been found in normal tissue, especially in the subcortical white matter and glial cells(*27*). Moreover, *TERT* promoter mutations have been detected as low-level somatic mosaicism in normal SVZ tissues in cases of IDH-wildtype GBM and in the hippocampus of Alzheimer’s patients(*21, 22*). Our findings suggest that clonal, low-level somatic mosaicism of *IDH* mutations in GPCs, including OPCs, of the normal cortex could predispose individuals to development of IDH-mutant gliomas. In support of this, a case study of two siblings demonstrated that somatic mosaicism of *IDH1^R132H^* acquired during early development in normal brain tissue and other regions predisposed them towards development of IDH-mutant astrocytoma(*67*).

Because of challenges in accessing various normal brain regions and the existence of potential sampling biases, the precise cell-of-origin for the initial driver mutation remains unclear in approximately 60% of our IDH-mutant glioma cohort. Nonetheless, our study provides direct evidence that GPCs (including OPCs) are the likely cells-of-origin for these tumors. This discovery not only sheds light on the evolution of IDH-mutant gliomas, but also suggests that these cells could be a source of somatic mosaicism linked to various neurological disorders(*68, 69*).

## Acknowledgments

The authors thank the patients and their families who took part in this study, the neurosurgeons who collected precious brain samples from the patients, and KAIST students, Do Hyeon Cha and Yeeun Seol, for their excellent technical assistance. We also acknowledge the facilities and the scientific and technical assistance of Dr. Youngsuk Hur of the EM & Histology Core Facility and Mrs. Jiye Kim at the FACS & NGS Core Facility of the BioMedical Research Center, KAIST.

## Funding

National Research Foundation of Korea funded by the Korean Government 2019R1A3B206661923 (JHL)

National Research Foundation of Korea funded by the Korean Government 2022R1A2B5B03001199 (S-GK)

MD-PhD/Medical Scientist Training Program through the Korea Health Industry Development Institute (KHIDI) funded by the Ministry of Health & Welfare, Republic of Korea (JWP)

Pioneer Research Center Program through the National Research Foundation of Korea funded by the Ministry of Science, ICT & Future Planning NRF2022M3C1A309202211 (S-GK)

Korea Health Technology R&D Project through the Korea Health Industry Development Institute (KHIDI), funded by the Ministry of Health & Welfare, Republic of Korea RS-2024-00438443 (S-GK)

Bio&Medical Technology Development Program of the National Research Foundation (NRF) funded by the Korean government (MSIT) RS-2024-00437820 (S-GK)

Sovargen (JHL)

## Author contributions

Conceptualization: JHL

Data curation: JWP, JK, SJ, J-HP

Formal analysis: JWP, JK, CHN, SHK, YSJ

Methodology: JWP, HJK, JHL

Resources: JY, J-KS, SP, AvD, JHC, S-GK

Investigation: JWP, K-WK, SML, CK, SA

Visualization: JWP

Funding acquisition: JWP, S-GK, JHL

Project administration: S-GK, JHL

Supervision: S-GK, JHL

Writing – original draft: JWP, JHL

Writing – review & editing: JWP, JHL

## Data and materials availability

Data from human whole-genome sequencing, 15 glioma-related gene panel sequencing, and mouse transcriptomes are deposited in the SRA (Sequence Read Archive) and GEO (Gene expression omnibus) with accession nos. PRJNA1149448 and GSE275791 are available for general research use. The human reference genome GRCh38 is available at https://www.ncbi.nlm.nih.gov/datasets/genome/GCF_000001405.26/, and the mouse reference genome GRCm39 is available at https://www.ncbi.nlm.nih.gov/datasets/genome/GCF_000001635.27/. In-house scripts for mouse transcriptomic analyses are available on GitHub (https://github.com/JeongHoLee-Lab/IDH-mutant_Glioma_cell_of_origin).

## Supplementary Materials

Materials and Methods

Figs. S1 to S17

Tables S1 to S5

References (*71–106*)

## Auxiliary Supplementary Materials

Tables S6 to S9

