## Supplementary material for "IDH-mutant gliomas arise from glial progenitor cells harboring the initial driver mutation": Materials and Methods

#### Collection of the IDH-mutant glioma patient cohort

Individuals with IDH-mutant glioma who had undergone supra-total resection since 2014 at Severance Hospital participated in this project. All patients were newly diagnosed with astrocytoma, IDH-mutant (*IDH1*$^{R132H}$), 1p/19q-non-codeleted; or oligodendroglioma, IDH-mutant (*IDH1*$^{R132H}$), and 1p/19q-codeleted; no patients had a previous history of surgery, chemotherapy, or radiotherapy (table S1). Standard peri-operative practices as described in previous reports were applied(*71*, *72*). Preoperative, high-resolution, axial magnetic resonance (MR) images – both contrast-enhanced T1-weighted and pre-contrast T2-weighted – were obtained for each patient on the day of surgery to localize the anatomical location. During the operation, tumor margins were checked using a neuro-navigation system, and matched normal cortex samples were collected on the way of surgical corridor (Fig. 1C). During supratotal resection of the tumor, the lateral ventricle was opened and SVZ specimens were carefully biopsied so as to preclude intraventricular hemorrhage. (Fig. 1C). All obtained normal cortex and SVZ samples were away from the tumor mass, with the precise location of the samples confirmed by the neuro-navigation system during surgery (Fig. 1B, and fig. S5). Three-dimensional distances between the biopsy point and its nearest tumor margin were measured using the US Food and Drug Administration (FDA)-approved medical imaging software, Aquarius intuitive edition 4.4.12 (TeraRecon), by postoperatively identifying these areas on preoperative T1 enhanced - (or T2) images (Fig. 1B, table S1, S4, and S5). Direct distances between tumor margin and normal cortex ranged from 3 to 25 mm (table S1, S4), whereas those between tumor margin and normal SVZ ranged from 0.8 to 5.02 mm (table S1, S5). After sample collection, microscopic tumor involvement was evaluated by histological examination according to WHO CNS5 criteria by an experienced neuropathologist(*2*). The histopathological examination included H&E staining; immunohistochemical (IHC) staining for the glioma marker proteins, IDH1$^{R132H}$, Ki67, p53 and ATRX; and fluorescence *in situ* hybridization (FISH)-based detection of 1p/19q-codeletion status. *O*$^{6}$-methylguanine DNA methyltransferase (*MGMT*) promoter methylation status was determined by methylation-specific PCR analysis of the *MGMT* gene promoter. Appropriate sampling and tumor-free status was also confirmed by H&E staining and IHC of cortical and SVZ tissues (Fig. 1B). All cases in this cohort were collected with informed consent according to protocols approved by the Institutional Review Boards of Severance Hospital (approval no. 4-2021-1319) and KAIST (approval no. KH2024-068), as well as the Committee on Human Research.

#### Collection of the IDH-negative control cohort

Cortex tissues from individuals of various ages with Lennox-Gastaut syndrome, focal cortical dysplasia type 2, polymicrogyria, hippocampal sclerosis, autism spectrum disorder, meningioma, metastatic brain tumor, ganglioglioma, or IDH-wildtype GBM were obtained for the control cohort of this project and confirmed as normal by an experienced neuropathologist (table S2). The biopsy protocol for other types of disease was similar to that for IDH-mutant glioma. Tissues from post-mortem human brains of Alzheimer disease and autism spectrum disorder patients were obtained from the Netherlands Brain Bank (Meibergdreef 47, 1105 BA

Amsterdam; Project ID: Lee-835) and the National Institute of Child Health and Human Development (NICHD). All cases in this cohort were collected with official consent from all subjects and approved by the Institutional Review Boards of KAIST (approval no. KH2024-68).

### DNA extraction from tissue samples

Genomic DNA was extracted from blood and brain specimens, including normal SVZ and normal cortex, tumor, and nuclei sorted by fluorescence-activated nuclei sorting (FANS). DNA was extracted from blood using a QIAamp DNA Blood Mini Kit (Qiagen), from bulk brain tissues using a QIAamp DNA Mini Kit (Qiagen), and from FANS-sorted nuclei using a QIAamp DNA Micro Kit (Qiagen), according to the manufacturer's instructions. The concentration of all extracted DNA was determined using an Invitrogen Qubit 4 Fluorometer (Thermo Fisher Scientific Inc.).

### Library preparation and hybrid capture sequencing of 15 glioma-related genes

Samples for hybrid capture sequencing of 15 glioma-related genes were prepared from 1 μg of extracted genomic DNA. Fifteen genes related with glioma in the TCGA database were selected (table S3)(*73–77*). Probes that effectively captured these 15 glioma-related genes were designed and manufactured by Celemics, Inc. Hybrid capture libraries were prepared according to the manufacturer's protocol and sequenced on a Miseq Dx platform (Illumina) by Sovargen, Inc. The average read depth of hybrid capture panel sequencing for these 15 genes was 723.1× in cortex samples, 679.9× in SVZ samples, and 880× in tumor samples.

### Library preparation and WGS

Samples for WGS were prepared from 1 μg of extracted genomic DNA using an Illumina TruSeq DNA PCR-Free Library Prep Kit (Illumina) and sequenced on NovaSeq 6000 and NovaSeq X plus platforms (Illumina) by Inocras Inc. WGS was performed on tumors and their matched samples, including normal cortex or blood, with an average read depth of 32.8×.

### Variant calling for gene panel sequencing and WGS

Analysis-ready bam files from sequenced reads were generated according to the GATK Best Practices workflow suggested by the Broad Institute. The fastq files were mapped to the human reference genome (GRCh38) using the Burrows-Wheeler aligner (BWA)-MEM algorithm. After the removal of duplicated reads by Picard, a matched sample analysis was performed using MuTect2(*78*), Strelka2(*79*) and Sequenza (used only for WGS)(*80*), which are specifically designed to analyze matched brain specimen–blood sample sequencing datasets. Single-nucleotide variants (SNVs), short indels, and copy-number variants (CNVs) were called as previously reported(*81*, *82*). False-positive events were removed by manually inspecting all alterations using the Integrative Genomics Viewer (IGV). Only variants with at least two supporting reads were accepted, and all panel sequencing and WGS results were validated by cross-checking with clinical sequencing data.

### Estimation of driver mutation timing

The WGS dataset used in this analysis, with a minimum average coverage of 28×, was derived from 20 patients in the PCAWG database(*26*), 12 in the TCGA database(*83*), and 6 in our IDH-mutant glioma patient cohort. Of these 38 IDH-mutant gliomas, 21 were diagnosed as astrocytoma and 17 as oligodendroglioma. The order and relative timing of driver mutations in IDH-mutant gliomas was estimated using the PhylogicNDT package with default parameters(*84*). Briefly, the Clustering submodule grouped the SNVs identified from WGS based on tumor cell fractions. Subsequently, the order and relative timing of driver mutations and copy number variations for each patient were estimated using the SinglePatientTiming submodule. Finally, the LeagueModel submodule integrated the timing of mutations across all patients within the cohort to infer the average order of mutations and estimate the odds ratio of mutations accumulating in the early (first half) or late (second half) stages of tumor evolution.

### Deep targeted amplicon sequencing and variant analysis

Deep targeted amplicon sequencing, used for validation sequencing, was performed on all candidate variants from each glioma patient, as previously described(*19*, *81*). Briefly, region-specific primers, with sequences shown in table S6, were designed using the Illumina Nextera single index (Illumina). Target sequences were amplified by PCR using PrimeSTAR GXL (#R050A, Takara) high-fidelity DNA polymerase under optimal thermal conditions. In a second amplification, up to 20 ng of purified product from the first PCR was annealed with both Illumina adaptor and index sequences. The final targeted amplicon sequencing libraries were pooled and sequenced on a Miseq Dx platform (Illumina) by Sovargen, Inc., with an average read depth 391,958×. Sequenced reads were sorted by index and generated fastq files, which were aligned to the human reference genome (GRCh38) using the BWA-MEM algorithm and then converted to bam files using Picard. A curated set of SNVs and short indels was prepared by manually inspecting all alterations with the IGV, as described above in panel sequencing and WGS analysis. In addition to manual inspection, an in-house algorithm implemented with Python was used to calculate allele frequency for each indel variant. Specifically, the algorithm determined whether each sequencing read supported the reference allele, an alternative allele, or neither. The algorithm simultaneously applies a filter to each sequencing read, screening out all but high-quality reads free of artifactual features. One read with more than two mismatched bases was removed during the filtering process. Thus, variant allele frequency for short indels, as well as the number of supporting reads for each allele, were calculated so as to only include high-quality reads that satisfied filtering criteria.

### Droplet-based digital PCR assays

The ddPCR QX200 system (Bio-Rad Laboratories, Inc.) was used for $IDH1^{R132H}$ detection. $IDH1^{R132H}$ mutations were identified using mutation-specific primer and probe combinations (BR186dHsaMDV2010055, Bio-Rad Laboratories, Inc.). Mutant and wildtype alleles were labeled with FAM and HEX, respectively. All ddPCR reactions included 5U of *Hae*III restriction enzyme (#R0108S, NEB) and were performed using the ddPCR Supermix for probes (#1863023, Bio-Rad Laboratories, Inc.) according to the manufacturer's protocol. Droplets were generated using the Automatic Droplet Generator QX200 (Bio-Rad Laboratories, Inc.) and analyzed with the QX200 Droplet Reader (Bio-Rad Laboratories, Inc.) according to the manufacturer's instructions. QuantaSoft analysis software (version 1.7.4; Bio-Rad Laboratories, Inc.) was used for processing data. To ensure sufficient sensitivity, we accepted only assays with > 100 copies/µl of wildtype (HEX+) DNA for further analysis. VAF for each sample was calculated as the fractional abundance of mutant (FAM+) relative to total (wildtype [HEX+] + mutant [FAM+]) DNA copies using QuantaSoft.

**Detecting and defining ultra-low-level (<1% of VAF) somatic mutations**

Discriminating between false-positive (noise) and true signals is important for detecting mutations in low mutational-burden samples. For ddPCR analysis, we checked the VAF of $IDH1^{R132H}$ of 33 normal cortices from the IDH-negative control cohort (table S2), from which we deduced the normal distribution curve of VAF of $IDH1^{R132H}$ (fig. S1B). Next, we calculated a Z-score (z) for the $IDH1^{R132H}$ mutational burden (VAF) of normal cortex in the IDH-negative control cohort according to the formula,

$$z = \frac{x-u}{\sigma}.$$

The Z-score showed that a cortical tissue carrying more than 0.115% of $IDH1^{R132H}$ VAF is statistically significant (greater than three standard deviations from the mean; Z-score > 3.0; $P < 0.00135$), indicating that $IDH1^{R132H}$ VAF values < 0.115% are considered false-positive signals. On the basis of this, we defined brain tissue from the IDH-mutant glioma cohort showing $IDH1^{R132H}$ VAF value > 0.115% from ddPCR analysis as *IDH1*-mutation–carrying tissue.

To increase the accuracy of our deep targeted amplicon sequencing data, we utilized RePlow(*28*), a computational method for detecting low VAF (<1%) somatic mutations, for deep targeted amplicon sequencing. This probabilistic model infers patterns of background errors (noise) in given amplicon sequencing data, and calls variants based on *P*-value (a *P*-value < 0.05 was used for accepting variants).

**Fluorescence *in situ* hybridization (FISH) for qualitative detection of 1p/19q-codeletion**

FISH was performed on 2-µm-thick sections of FFPE specimens of three IDH-mutant gliomas (G5, G19, and G25) and their matching normal cortices. To accurately access the codeletion of chromosome arms 1p/19q in the IDH1-shared group cortex, we used the G5 and G19 tumor specimens (diagnosed as OD with 1p/19q-codeletion) as positive controls. Tumor and normal cortex specimens of G25 (diagnosed as AS with intact 1p/19q) were used as negative controls. Only two normal cortices from the IDH1-shared group (G5, G19) were utilized to evaluate 1p/19q status. Two slides from each specimen were prepared: one for detection of 1p status, and the other for detection of 19q status. The remainder of the experimental process followed the manufacturers' instructions. Briefly, slides were deparaffinized and incubated with pretreatment solution at 80°C for 60 minutes, then digested with protease solution at 37°C for 60 minutes. After the fixation process, samples were stained by dual-color-probe hybridization (#Z-2272-20, ZytoVision GmbH), with red probes labeling the test regions, 1p36 and 19q13, in the corresponding chromosome, and green probes labeling the control regions, 1q25 and 19p13, in the corresponding chromosome. Nuclei were counterstained with Leica mounting solution containing 4',6-diamidino-2-phenylindole (DAPI).

To increase the sensitivity of FISH, which has a limited ability to detect low-level somatic mosaicism, we counted a total of 400 non-overlapping nuclei for 1p and 19q (compared with the 100-120 non-overlapping nuclei usually counted in standard protocols(*85*, *86*)) in both normal cortices in the IDH1-shared group. FISH images were interpreted by an experienced neuropathologist according to previously reported guidelines(*87*). Slides in which the test/control (red/green) ratio was less than 0.8 were considered positive for 1p/19q-codeletion.

**Analysis of tumor-restricted mutations**

Starting with vcf files from matched WGS data from four individuals (G11, G20, G21, and G35), we randomly selected 9 to 18 tumor-restricted alterations, including driver mutations, in each case. All selected tumor-restricted mutations were SNVs or short indels. VAF of tumor-restricted mutations in tumor tissue ranged from 11.62% to 75.4% (except for the *RB1* mutation in G11, with a VAF of 0.2%). The variant effect predictor (VEP)(*88*) of all tumor-restricted passenger variants was "Low" or "Modifier". Shared mutations in normal cortex were then verified by performing deep targeted amplicon sequencing. Subsequent analyses followed the previously described method.

**Nuclei sorting from frozen human brain specimens**

After initially homogenizing tissue with 1% (wt/vol) formaldehyde in Dulbecco's phosphate-buffered saline (DPBS) without $Ca^{2+}/Mg^{2+}$, brain nuclei were sorted as previously described(*89*), then stained overnight with NEUN Alexa Fluor 488 antibody (clone A60, 1:2,500; #MAB377X, Millipore) and OLIG2-Alexa Fluor 647 antibody (clone EPR2673, 1:2,500; #81886S, Abcam). Nuclei were washed the following day with 4 ml FANS buffer, passed through a 35-µm cell strainer snap cap, and stained with 0.5 µg/ml DAPI (#422801, BioLegend). Stained nuclei were isolated with a FACSAria Fusion (BD) or MoFlo Astrio EQ cell sorter (Beckman Coulter) and pelleted at 1,600 × *g* for 5 minutes at 4°C in FACS buffer. Isolated nuclei were immediately processed for DNA extraction. Nuclei sorting experiments were carried out in the FACS Core Facility at the BioMedical Research Center, KAIST, and the Research Solution Center at the Institute for Basic Science.

**Management of transgenic mice**

All animal experiments were approved by the Institutional Animal Care and Use Committee (IACUC) of KAIST, and all experiments were performed according to IACUC guidelines. LSL-Cas9 (The Jackson Laboratory, #026175)(*90*) and $Idh1^{fl(R132H)}$ (a kind gift from Dr. A. Von Deimling)(*32*) mice on a C57BL/6 strain background were housed in isolator cages with free access to food and water and maintained under specific-pathogen-free conditions at a constant temperature of 23°C on a 12-h light-dark cycle (lights off at 19:00). The health status of mice was examined regularly by veterinarians and investigators. Disease-specific survival end points were death of mice or satisfaction of criteria for euthanasia under the IACUC protocol. The criteria for euthanasia were (i) severe weight loss (>20%); (ii) severe neurological impairment, including paralysis, seizure, and hunched posture with impaired motor power; or (iii) signs of head bulging.

### Post-natal *in vivo* electroporation

Neonatal 0–1-day-old pups (P0–1) were anesthetized by hypothermia (>5 minutes) and fixed to a support using an adhesive plaster. A virtual line connecting the right eye with lambda was used as a general positional marker, and a capillary needle was inserted at about one-third the length of this line from the eye. The right lateral ventricle was injected at a depth of 1 mm from the surface with 1 μl of plasmid mixture solution (>2 μg/μl, containing 1% Fast Green). All plasmid solutions were mixed with the donor vector and PBase-expressing helper vector (Donor:Helper molar ratio, 2:1). Injection success was verified by visualizing the shape of the Fast-Green–stained lateral ventricle, and only successfully injected animals were subjected to electroporation. For electroporation, animals were subjected to five electrical pulses (100V, 50 ms) at 950-ms intervals using an ECM830 electroporator (BTX-Harvard Apparatus) and 1-mm tweezer electrodes (#CUY650P1, Nepagene). The positive electrode was positioned ahead of the eye, and the negative electrode was placed in the opposite position on the ventral side. After electroporation, mice were placed on a 37°C-heating plate until they fully recovered and then were returned to their mothers.

### Validation sequencing of altered *Idh1* (*Idh1*$^{R132H}$), *Atrx*, and *Cdkn2a* in mouse samples

The edited genome status of our targeted genes, *Idh1*, *Atrx* and *Cdkn2a* was confirmed by isolating NSCs in the SVZ from a transfected mouse and culturing them as neurospheres *in vitro* using a modification of previous methods for FACS-based isolation of human-derived neural stem cells(*94*). In brief, fresh mouse SVZ specimens were minced with a scalpel in PIPES (#P1851-100G, Sigma-Aldrich) solution, then incubated in papain digestion solution (#LS003120, Worthington Biochemical) containing DNase I (#LS002139, Worthington Biochemical) at 37°C for 15 minutes. After centrifuging the digested tissue, the enzymatic digestion was blocked with a papain/trypsin inhibitor in DMEM/F-12 (#LM002-08, Welgene) media. Next, single-cell suspension media plus growth factors – 20 ng/ml of basic fibroblast growth factor (bFGF; $236-EG-01M, R&D Systems), 20 ng/ml of epidermal growth factor (EGF; #233-FB-025, R&D systems), and 50 U/ml penicillin/50 mg/ml streptomycin – was distributed in 6-well, low-attachment plates (#3471, Corning) and cells were incubated for 1 week, with media changes every 2 days. On day 7, cells were collected based on their GFP status and cultured for 1 additional week with a media change. The collected GFP-positive and -negative neurospheres were then progressively expanded.

For subsequent Sanger sequencing and amplicon sequencing, genomic DNA from GFP-positive and -negative neurospheres was extracted using the QIAamp micro DNA kit (Qiagen) following the manufacturer's instructions. Sanger sequencing of the mouse *Idh1* region was performed by Bionics, Co. Ltd. Targeted amplicon sequencing of the *Atrx* and *Cdkn2a* area was conducted as described above ("Deep targeted amplicon sequencing and variant analysis"). The primers for validation of mouse genome editing are listed in table S6. The efficiency of the sgRNAs, sgTrp53 and sgNf1, were confirmed in a previous report(*33*).

#### Stereotaxic injection of AAV5 into the cortex

Adeno-associated viruses (AAV5) targeting mature astrocytes, neurons, and oligodendrocytes were generating by a single viral vector containing sgRNAs targeting *Trp53*, *Atrx*, *Nf1* and Cre recombinase under control of the CBh promoter, and packaging it using VectorBuilder (VectorBuilder ID, VB230505-1625frj).

For stereotaxic injections, 4-week-old mice were anesthetized by inhalation of 5% isoflurane (Piramal healthcare) in an air/$O_2$ mixture and positioned in a stereotaxic frame (Stoelting Co.). Injections were performed unilaterally using a capillary needle containing 100 nl of virus solution ($1 \times 10^{13}$ to $1 \times 10^{14}$ genome copies/ml) at a flow rate of 25 nl/min using a syringe pump (#87930, Hamilton Company). The cortex was infected with a small amount of virus to prevent targeting of many OPCs. Discharge along the injection track was prevented by leaving the needle in place for 3 minutes after the injection and then slowly withdrawing it. Coordinates for injections were -1, 1.7, and 1 mm caudal, lateral, and ventral relative to bregma. After the injection, mice were placed on a 37°C heating plate until they fully recovered and then were returned to their cages.

#### Image analysis of mouse brain sections

The brains of treated mice were fixed in freshly prepared phosphate-buffered 4% paraformaldehyde, cryoprotected overnight in 30% buffered sucrose, formed into blocks of optimal cutting temperature compound (#3801480, Leica Biosystems), and stored at -80°C. Cryostat-cut sections (40 µm thick) were collected and stored in 50% (vol/vol) glycerol in phosphate-buffered solution at -20°C. For H&E and DAB (3,3'-diaminobenzidine) staining, formalin-fixed, paraffin-embedded sections were cut at a thickness of 4 µm. The collected brain sections were stained in a free-floating manner with the following antibodies: chicken anti-GFP antibody (1:1,000 dilution; #AB13970, Abcam), rabbit anti-OLIG2 antibody (1:500 dilution; #AB9610, Millipore), rabbit anti-NG2 antibody (1:500 dilution; #AB5320, Merck), mouse anti-NEUN antibody (clone 60; 1:500 dilution; #MAB377, Millipore), rabbit anti-GFAP antibody (1:500 dilution; #z0334, DAKO), mouse anti-adenomatous polyposis coli (APC) antibody (clone CC-1; 1:50 dilution at 4°C for 10 days; #OP80, Millipore), mouse antibody to IDH1$^{R132H}$ (1:20 dilution after heat-induced epitope retrieval; #DIA-H09, Dianova), mouse anti-ATRX (D-5 clone; 1:500 dilution; #sc-55584, Santa Cruz), rat anti-Ki-67 antibody (1:500 dilution; #14-5698-82, eBioscience), and mouse anti-Nestin antibody (1:500 dilution; #MAB353, Sigma). DAPI, included in mounting solution (#P36931, Life Technology), was used for nuclear staining. Images were acquired using a Zeiss LSM980 confocal microscope and Axio scan Z1 (Carl Zeiss). Slide scanning experiments were carried out in the EM & Histology Core Facility at the BioMedical Research Center, KAIST, and in the Research Solution Center at the Institute for Basic Science. Fluorescence intensities reflecting the distribution of GFP-positive cells were converted to gray-scale values and measured using ImageJ software (https://imagej.net/ij/), which was also used to count number of GFP-positive cells in virus-injected mice.

#### Library preparation for scRNA-seq analysis of mouse IDH-mutant glioma model

Single-cell isolation was performed following the manufacturers' instructions. Briefly, four IDH-mutant glioma model mice (OPC-ITAN, 26 to 38 weeks old) and a control mouse (OPC-LacZ,

20 weeks old) were euthanized under isoflurane anesthesia by transcardiac perfusion of 20 ml cold DPBS without $Ca^{2+}/Mg^{2+}$. Brains were harvested and washed in DPBS without $Ca^{2+}/Mg^{2+}$. GFP-positive areas in the brain were micro-dissected with a scalpel blade under an inspection lamp (#UV-BL-FL5000, NDT Supply.com, Inc.), pooled for digestion using the Adult Brain Dissociation Kit, mouse and rat (#130-107-677, Miltenyi Biotec), and kept on ice. After manual, gentle tissue dissociation, cell debris in the single-cell suspension was removed by gradient centrifugation (3000 × g for 10 minutes) at 4°C. Single-cell suspensions were labeled with antibodies against immune cell markers – CD11b-APC (1:100 dilution; #101212, BioLegend) and CD45-APC-Cy7 (1:100 dilution; #103116, BioLegend) – in flow cytometry buffer and counterstained with DAPI (0.5 µg/ml; #422801, BioLegend) to discriminate dead cells from live cells. Stained single-cell suspensions were isolated using a FACSAria Fusion (BD). The resulting $DAPI^-$/$immune^-$/$GFP^+$ and $DAPI^-$/$immune^-$/$GFP^-$ cells were used for subsequent 10X scRNA-seq analysis. About 7,500–10,000 live cells from each of our five mice were labeled with a CellPlex kit (#PN-1000261, 10X Genomics), and a scRNA-seq library was prepared using Chromium Single Cell 3' kit v3.1 (#PN-1000268, 10X Genomics) according to the manufacturer's protocols. Single-cell sorting and library preparation were carried out in the FACS & NGS Core Facility at the BioMedical Research Center, KAIST, and the library was sequenced on the NovaSeq X plus platform (Illumina) by Inocras Inc.

**scRNA-seq data processing**

scRNA-seq raw sequencing data were processed using Cell Ranger (v.8.0.0). Samples were aligned to a custom transcriptome reference based on the mouse reference genome, GRCm39 (with the Cas9-EGFP sequence added), using Cell Ranger mkrefs. One of the five samples was processed using the Cellranger count workflow, whereas the other multiplexed samples were processed using Cellranger multi. Ambient RNA was removed with Cellbender v0.3.1(*95*) using the raw output from Cell Ranger, with expected-cells set to 10,000 for the single sample and 20,000 for multiplexed samples. Additional data analysis was carried out using Python 3.10 and Scanpy (v1.9.8), based on published workflows(*96*). For plotting, the mplscience package (v0.0.7, https://github.com/adamgayoso/mplscience) was used for aesthetic visualization.

**scRNA-seq preprocessing**

Low-quality cells with fewer than 200 genes in filtered outputs of the Cellbender pipeline were removed. Additional quality control was performed using 5 median absolute deviations (MADs) of log1p_total_counts, log1p_n_genes_by_counts, and pct_counts_in_top_20_genes, and 3 upper MADs of pct_counts_mt. Doublets were also removed using scDblFinder (v1.16.0). The top 3000 highly variable genes were selected using the Seurat v3 method, and mouse samples were batch-corrected and integrated using scVI(*97*) with hyperparameter tuning ('n_hidden' = 192, 'n_latent' = 60, 'n_layers' = 1, gene_likelihood = 'zinb', dropout_rate = 0.5, with mitochondrial & ribosomal percentages as covariates). Normalization was then performed using Scanpy's sc.pp.normalize_total() and sc.pp.log1p(). Neighbors were calculated using the latent representation of the scVI model and clustered using the Leiden algorithm. Cell type annotation was performed by manual annotation of marker genes using the markers in table S8. This process was assisted by Celltypist (v1.6.2)(*98*) mapping to a normal mouse brain atlas and marker gene-based annotation using CellID (v1.21)(*99*).

#### Non-negative matrix factorization and gene expression module assignment

Gene expression modules in tumor-lineage cells (OPCs, COPs, and tumor cells) were identified by subclustering tumor-lineage cells and filtering out genes expressed in fewer than 10% of cells. Subclustered samples were re-integrated using scVI, setting mitochondrial and ribosomal percentages as covariates. The cNMF package (v1.5.4)(*41*), with downstream analysis scripts kindly shared by Dylan Kotliar, was utilized after model training with all 25,231 genes instead of only highly variable genes. In addition to the default parameters in the cNMF pipeline, we attempted to correct for batch effects using TPM-normalized expression from the fully trained scVI model and re-selecting the top 2000 highly variable genes in tumor-lineage cells. NMF performance was compared using 300 iterations and a range of metamodules (K) from 2 to 10. From the diagnostic plot, we chose K = 8 for downstream analysis (fig. S15A, marked with green arrow). Gene expression programs from cNMF output were named using known marker genes(*7*, *8*, *38*).

#### Comparison with transcriptomes of human brain tumors

Transcriptomic profiles of mouse IDH-mutant glioma models and different human brain tumors (OD, AS, and GBM) were compared using publicly available datasets from previous publications(*38*, *43*). Mousipy (v0.1.5, https://github.com/stefanpeidli/mousipy) was used to translate human gene symbols to mouse gene symbols from publicly available datasets. For each batch, scRNA-seq data were pseudobulked into one sample, then normalized using sc.pp.normalize_total() and sc.pp.log1p(). For the mouse tumor model, each sample was pseudobulked using three pseudo-replicates for better comparisons. Pairwise distances and correlations were calculated using scikit-learn (v1.4.1) and plotted with the seaborn clustermap.

#### Pseudotime and trajectory analysis

Pseudotime inference and trajectory analyses were performed using Monocle3 v1.3.7(*100*) with R 4.3.3. Re-calculated UMAP embeddings obtained using previous scVI embeddings and the expression matrix from tumor-lineage subcluster cells were transferred to Seurat (v5.1.0) and normalized using SCTransform(*101*). The Seurat object was then transformed into a SingleCellExperiment object using Monocle's default pipeline. Root cells were selected among OPCs, with introduction of driver mutations using pr_graph_cell_proj_closest_vertex instead of manual selection of specific root cells. Calculated pseudotime was extracted, and normalized gene expression modules per cell from NMF results were plotted against pseudotime by fitting with a generalized additive model from pygam (v0.9.1)(*102*) and applying multiple testing correction using the Benjamini & Hochberg method.

#### Differentially expressed gene (DEG) analysis

DEG analysis was performed using pyDESeq2 (v0.4.9)(*103*) with pseudobulk, which increases the robustness of the analysis by creating three pseudo-replicates per condition To identify the variance of gene expression attributable to the effects of driver mutations, we integrated normal OPCs from the adult mouse brain atlas(*44*). DEGs were compared across various conditions (tumor cells vs. GFP-positive OPCs from tumor model mice; GFP-positive OPCs from tumor model mice vs. OPCs from the public database). The Sanbomics package (https://github.com/mousepixels/sanbomics) was used for volcano plots, and functional analyses were performed using decouplR (v 1.4.0)(*104*) and gseapy (v1.13)(*105*) with Mouse Hallmark(*70*).

**Statistical analysis**

Data are presented as means ± standard error of mean (s.e.m.). Results were analyzed by *t*-test, analysis of variance (ANOVA) or Fisher's exact test, as appropriate, using GraphPad Prism version 9 (GraphPad Software, Inc.). Assuming normal distribution curve of $IDH1^{R132H}$ in IDH-negative control cohort, we utilized Origin2019 (OriginLab Corporation). ComplexHeatmap package (ver 2.18.0)(*106*) was utilized for summarizing sequencing data from our patient cohort. A Kaplan-Meier analysis was applied to survival data for mice. *P*-values less than 0.05 were considered statistically significant, except for finding the *IDH1* mutation carrying tissue, where a *P*-value less than 0.00135 was used. All animals used in experiments or as a source of cells were subjected to randomization. Sample sizes were predetermined based on the variability found in preliminary and similar experiments. Researchers were not blinded to allocation during experiments or analysis of outcomes.
